# Hydraulic control of embryo size, tissue shape and cell fate

**DOI:** 10.1101/389619

**Authors:** Chii Jou Chan, Maria Costanzo, Teresa Ruiz-Herrero, Gregor Mönke, Ryan J. Petrie, L. Mahadevan, Takashi Hiiragi

## Abstract

Size control is fundamental in tissue development and homeostasis^1,2^. While the role of cell proliferation in this process has been widely studied^3^, the mechanisms of organ size control and how it impacts cell fates remain elusive. Here, we use mouse blastocyst development as a model to unravel a key role of fluid-filled lumen in embryonic size control and cell fate specification. We find that during blastocyst expansion, there is a two-fold increase in the pressure of the lumen that translates into a concomitant increase in the cortical tension of trophectoderm (TE) cells lining the lumen. Increased cortical tension leads to vinculin mechanosensing and maturation of the functional tight junctions, thereby establishing a positive feedback loop to accommodate lumenal growth. However, when the cortical tension reaches a critical threshold, cell-cell adhesion cannot be sustained, and mitotic entry leads to a rupture of TE epithelium, fluid leakage and collapse of the blastocyst cavity. A simple theory of hydraulically-gated oscillations that integrates these feedback interactions recapitulates the evolution of cavity size and predicts the scaling of embryonic size with the tissue volume. Our theory further predicts that reduced cortical tension or disrupted tight junctions, and increased tissue stiffness lead to smaller embryonic size. These predictions are verified experimentally by embryological, pharmacological and genetic manipulations of the embryos. Remarkably, these changes to lumenal size, without a change in the tissue volume, lead to alteration of tissue architecture and cell fate. Overall, our study reveals how lumenal pressure and tissue mechanics control embryonic size at the tissue scale, that in turn couples to cell position and fate at the cellular scale.

---

Studies on organ size control generally focus on the regulation of cell proliferation. However, tissue size and pattern also depends on cell size, shape, and spatial organization, all of which can be influenced by tissue-generated mechanical forces. In particular, lumenal pressure is thought to play a central role in shaping early development of vasculature, renal system and endocrine organs^4^. Yet, our understanding of how lumenal pressure impacts organ size and shape, and how these are coordinated with cell fate remains primitive. To make headway on these questions requires a system that is both simple and capable of recapitulating important aspects of development. In this study, we use the mouse blastocyst as a model system to investigate these mechanisms in just such a minimal setting.

Blastocyst formation is a key process in early development that is conserved across mammalian species^5^. As the fluid-filled cavity forms and expands^6^, the cells in the embryo segregate into the extra-embryonic trophectoderm (TE) lining the cavity and the inner cell mass (ICM); the TE eventually gives rise to the placenta while part of the ICM develops into the embryo proper^7,8^. Following this process dynamically, we observed that initially the blastocyst cavity shows a steady increase in volume, but as it matures, this steady growth transitions to an intermittent mode wherein the blastocyst alternates between growth and sudden collapse; these events become more frequent towards the late blastocyst stage (E4.5) (Fig. 1a). Since these pre-implantation developmental processes occur within a stiff glycoprotein coat known as the zona pellucida (ZP), blastocyst size has generally been assumed to be limited by the ZP, although the ZP was also shown to be dispensable for mouse development^9^. Indeed, blastocysts without the ZP or even those that form after embryo dissociation and re-aggregation, display cycles of expansion and collapses until they reach a similar size to that seen in the late blastocyst stage with the ZP (Fig. 1b, Extended Data Fig. 1a, b). Furthermore, the dynamics of intermittent collapse, as measured by the time between collapses (Inter-collapse interval, ICI) for blastocysts with and without ZP is very similar (Fig. 1c), which suggests that the ZP does not influence the dynamics of blastocyst collapse and its ultimate size. Together, these observations point to an autonomous mechanism for blastocyst size control independent of the potential mechanical constraint imposed by the ZP.

**Figure 1:**
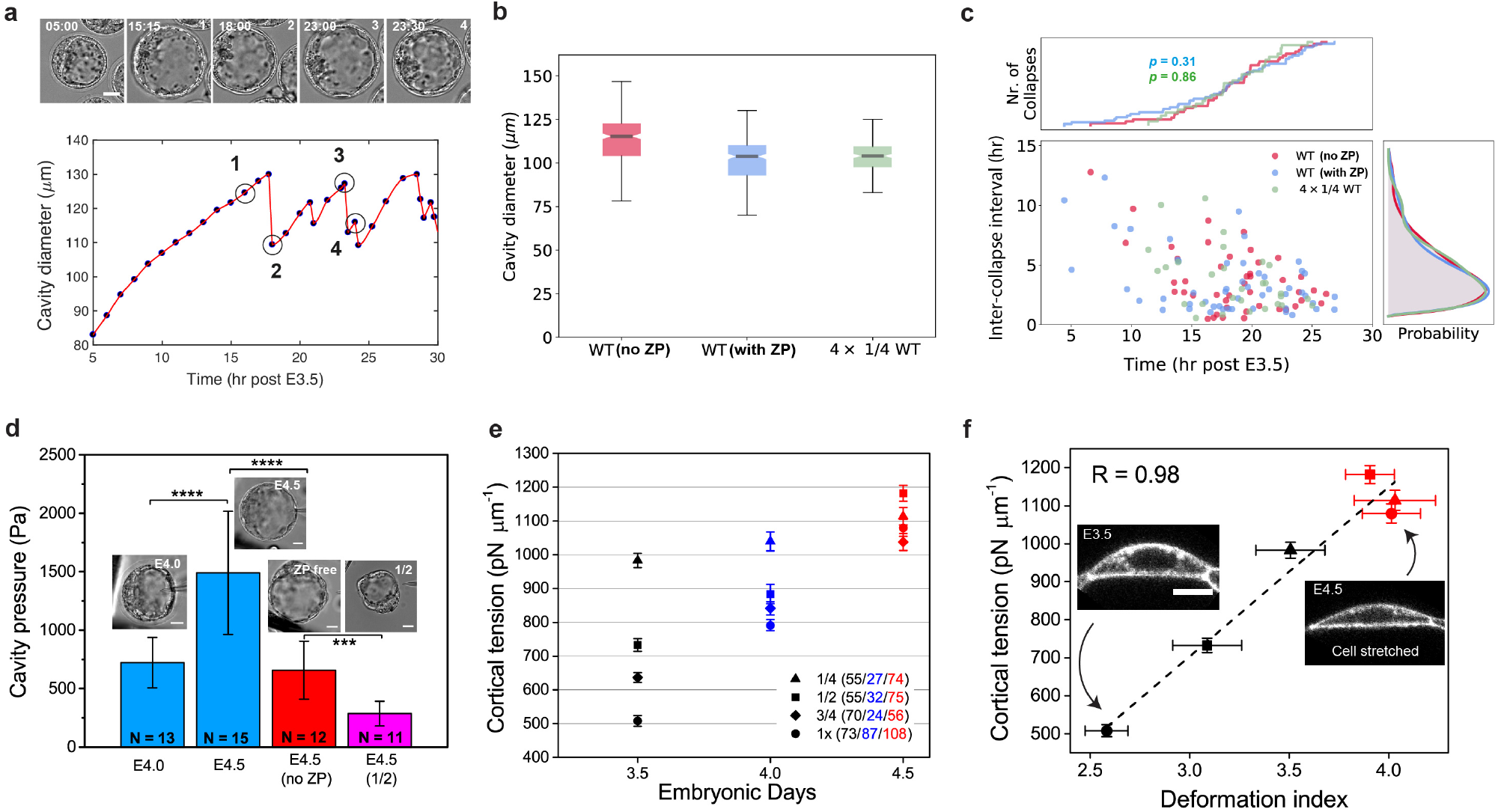
Blastocyst development is driven by an increase in lumenal pressure and TE cortical tension. **a.** Top panel: Representative images of a *zona pellucida* (ZP)-intact blastocyst undergoing blastocyst expansion. Time (hh:mm). Bottom panel: Corresponding plot of cavity diameter as a function of time (post E3.5). Time (hr) post E3.5 (84 hr post-hCG injection). Dotted line denotes the cavity. **b.** Cavity diameter at the plateau stage for wild-type (WT) blastocysts with (blue) and without (red) ZP, and for aggregated embryos composed of 4 × 1/4 embryos (green). c. Inter-collapse interval (ICI) plotted as a function of time (post E3.5), for WT with (blue, *N* = 13) and without (red, *N* = 14) ZP, and aggregated embryos composed of 4 × 1/4 embryos (green, *N* = 13). Kolmogorov-Smirnov test for the number of collapse events (cumulative distribution) compared to the WT (no ZP) embryos: *P* = 0.31 for WT (with ZP) embryos and 0.86 for 4 × 1/4 embryos, suggesting that the three distributions are not statistically different. **d.** Cavity pressure of full blastocysts at various developmental stages, with and without ZP. Pressure for 1/2 blastocyst at E4.5 is also depicted. *N* = number of blastocysts measured. Scale bars: 20 μm. **e.** Cortical tension of TE in developing blastocysts of various reduced systems, measured during early (E3.5, black), mid (E4.0, blue) and late blastocyst stage (E4.5, red). Numbers in the brackets indicate the number of cells measured in each state/stage. **f.** Cortical tension as a function of deformation index, defined as TE basal area normalized to the cell volume. Colors and symbols correspond to those in (e). *N* (E3.5) = 36, 45 and 46 for 1/4, 1/2 and 1× embryos, respectively. *N* (E4.5) = 53, 58 and 49 for 1/4, 1/2 and 1× embryos, respectively. Error bars = SD. Scale bar: 20 μm. \*\*\*\**P*< 0.0001, \*\**P*< 0.01

Given that the blastocyst is a fluid-filled multicellular tissue shell, we turn to a consideration of the lumenal pressure and the cortical tension (CT) of the TE epithelium as key regulators of blastocyst size. Using a micropressure probe^10^, we directly measured the hydrostatic pressure *in vivo* in a developing mouse embryo (Extended Data Figure 2). Midstage blastocysts (E4.0) have a lumenal pressure of approximately 700 Pa, which increases two-fold to more than 1500 Pa by the mature blastocyst stage (E4.5; Fig. 1d). In contrast, blastocysts without the ZP show a lower pressure (Fig. 1d, red), indicating additional pressure build-up due to the constricting nature of the ZP. When the blastocyst tissue volume is embryologically manipulated by bisecting the 4-cell stage embryo, the resulting E4.5 half blastocyst has a smaller size and reduced pressure. Independently, the CT of TE cells, measured by micropipette aspiration^11^, shows a progressive increase during blastocyst development. At early blastocyst stage (E3.5), the TE cells in reduced-size blastocysts (1/4, 1/2 and 3/4) exhibit higher CT than the whole embryos. However, as the blastocyst matures and reaches its steady-state size at E4.5, the CT of all blastocysts converges to a similar critical value (~1100 pN/μm) (Fig. 1e). These observations that the lumenal pressure increases during blastocyst development, and that the final CT converges to the same critical value independent of blastocyst size, suggest that hydraulically generated tissue tension regulates mature blastocyst size.

Moving from the tissue to the cellular scale, we next quantified cell shape changes across all blastocysts at various developmental stages. We found that the CT scales remarkably with cell deformation (Fig. 1f, Extended Data Figure 1c, d), which suggests that the lumenal pressure indirectly affects TE cell mechanics globally, via cell stretching, in a cell non-autonomo^u^s manner. To differentiate correlation from causation between cavity expansion and cell geometry change, we first perturbed cavity expansion by pharmacologically blocking fluid influx or inducing fluid leakage. Impaired function of Na^+^/K^+^ ATPase pumps by ouabain^12^ or of tight junction integral membrane protein claudins by *Clostridium perfringens* enterotoxin (CPE)^13^ led to reduced cavity size and CT of TE cells (Fig. 2a). We next mechanically inflated (deflated) the cavity on short timescales (~30 s) using a microinjection pipette, and observed a transient increase (decrease) in the CT of TE cells (Fig. 2b). These data consistently indicate that the cavity expansion drives changes in tissue and cell geometry, which in turn increases the TE cortical tension.

**Figure 2:**
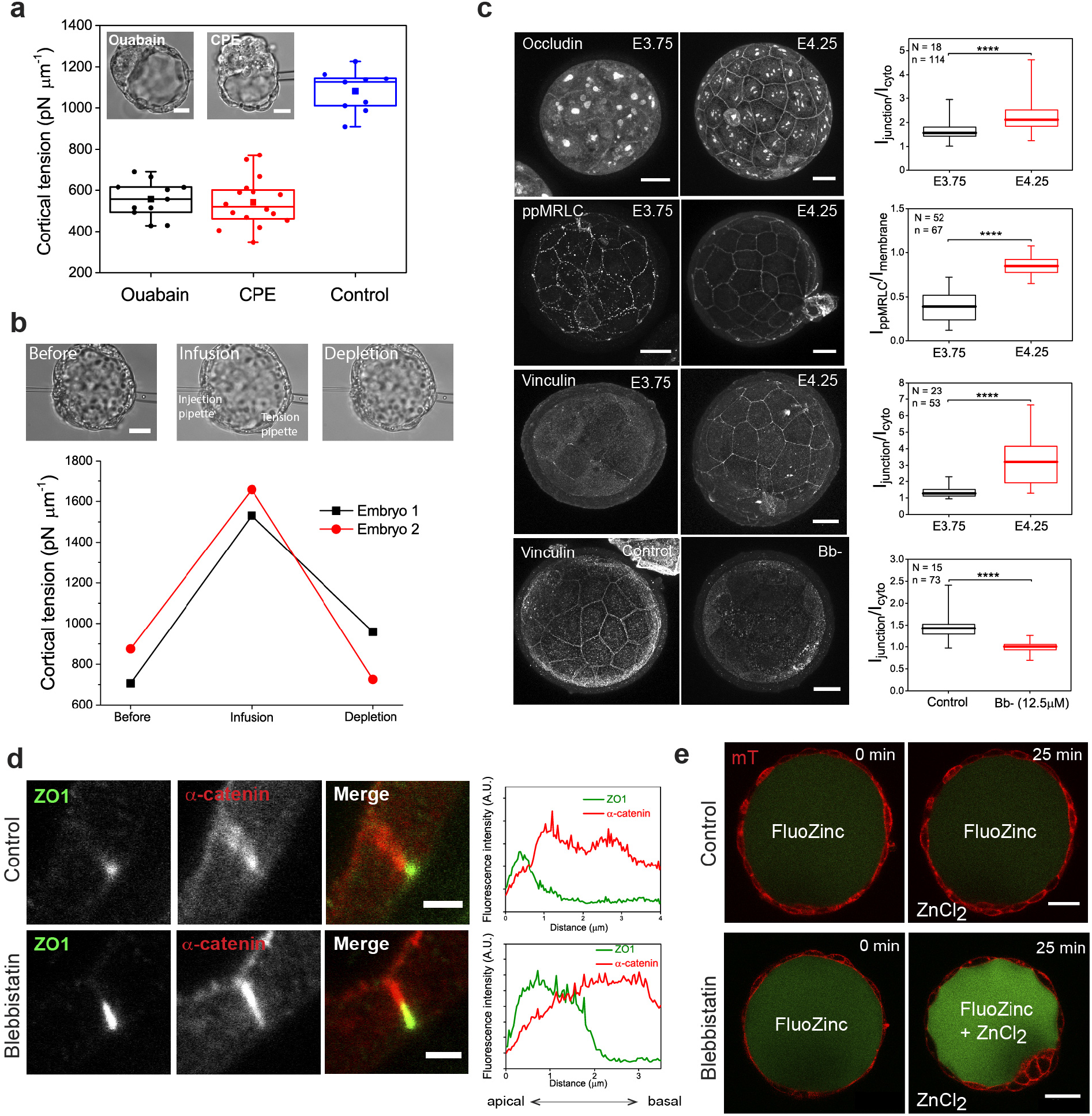
Lumenal expansion promotes junctional maturation. **a.** Cortical tension of TE cells in embryos treated with 500 μM of ouabain (*n* = 14, *N* = 5) and 100 μg/ml of CPE (*n* = 16, *N* = 7) compared to the control embryos (*n* = 9, *N* = 4). Scale bars: 20 μm. **b**. Cortical tension of TE cells from two representative blastocysts, following mechanical infusion and depletion of cavity fluid at short timescales (~30 s). Scale bar: 20 μm. **c.** Immunostaining of occludin, di-phosphorylated myosin regulatory light chain (ppMRLC) and vinculin at early (E3.75) and late (E4.25) blastocyst stage, and of vinculin upon 12.5 μM Bb-treatment (E4.25). Right panels: corresponding box plots showing junctional recruitment of these proteins during blastocyst expansion. *N* = number of embryos and *n =* number of blastomeres. Scale bars: 20 μm. **d**. Immunostaining of ZO1 and α-catenin in control versus Bb-treated (12.5 μM) blastocysts (E4.25). Line scans for the respective conditions are shown on the right, with 0 μm indicating the apical-most part of the tight junction. A.U., arbitrary units. Scale bars: 20 μm. **e**. Tight junction leakage assay using 100 μM Fluozinc and 200 μM ZnCl_2_. Blastocysts loaded with Fluozinc and treated with control (Bb+) and ZnCl_2_ showed no increased signals within the blastocoel with time (top panels), in contrast to the enhanced signals within the blastocoel with Bb-treatment for the same duration (bottom panels). Scale bars: 30 μm. \*\*\*\**P*< 0.0001.

To understand the molecular basis for these events, we note that stretch-induced increase of CT is associated with a series of cytoskeletal remodeling events at TE cell-cell junctions (Fig. 2c). Specifically, we observed a change in the localization of occludin, a transmembrane tight junction protein, from cytoplasm to cell-cell junction, following blastocyst expansion. This is accompanied by an enrichment of di-phosphorylated myosin regulatory light chains (ppMRLC) at tight junctions (Extended Data Fig 3a), and a shift in localization from punctate to more continuous pattern (Fig. 2c). Vinculin, a tension sensor reported *in vitro^14^*, shows strong enrichment at the TE tight junctions following blastocyst maturation, and an expression level that correlates with the measured CT (Extended Data Fig. 3c-f). Conversely, either treatment with the myosin II inhibitor blebbistatin (Bb-) or mechanical deflation leads to the disassembly of vinculin and ZO1 from tight junctions (Fig. 2c, d, Extended Data Fig. 3g), indicating that junctional maturation is tension-dependent. Specific localisation of ppMRLC, Myh9 and vinculin at tight junctions (Extended Data Fig. 2a-d) suggests that tight junctions are likely the stress-bearing sites during blastocyst expansion, unlike other epithelia in culture or tissues where vinculin are localised at the adherens junctions^15–17^. In agreement with this, reducing CT by Bb-disrupted the tight junction seals, as revealed by FluoZinc/ZnCl_2_-based permeability assay (Fig. 2e). Together, these results show how a positive feedback loop linking tension and tight-junction maturation allows the blastocyst to accommodate high pressure build-up during cavity expansion.

In mature blastocysts, the presence of a constant final TE cortical tension independent of size above which the blastocysts collapse intermittently (Fig. 1e) implies the existence of a critical failure stress in the TE epithelium. Indeed, we observed that blastocyst collapse is often preceded by mitotic events (Fig. 3a), suggesting that the collapses may be triggered by junctional loss through active cell rounding. Previous studies have shown that mitotic cell rounding leads to increased CT^18^, which could transiently impair tight junction sealing and lead to blastocyst collapse. Live imaging of blastocysts loaded with FITC-dextran dye confirmed this by revealing fluid leakage through the cell-cell junctions involving mitotic cells (Extended Data Fig. 4a). To further test this hypothesis, we blocked cell cycle progression during blastocyst expansion. Embryos arrested at the S-phase by aphidicolin treatment showed almost no collapse events compared to the controls (Fig. 3b upper panel, Extended Data Fig. 4c). However, those treated with nocodazole and arrested at the M-phase showed constant fluid leakage and failed to expand (Fig. 3b lower panel, Extended Data Fig. 4d), which could be rescued by washout (Extended Data Fig. 4e). These findings support a model in which mitosis may trigger a temporal disintegration of TE junctional complex leading to blastocyst collapse.

**Figure 3:**
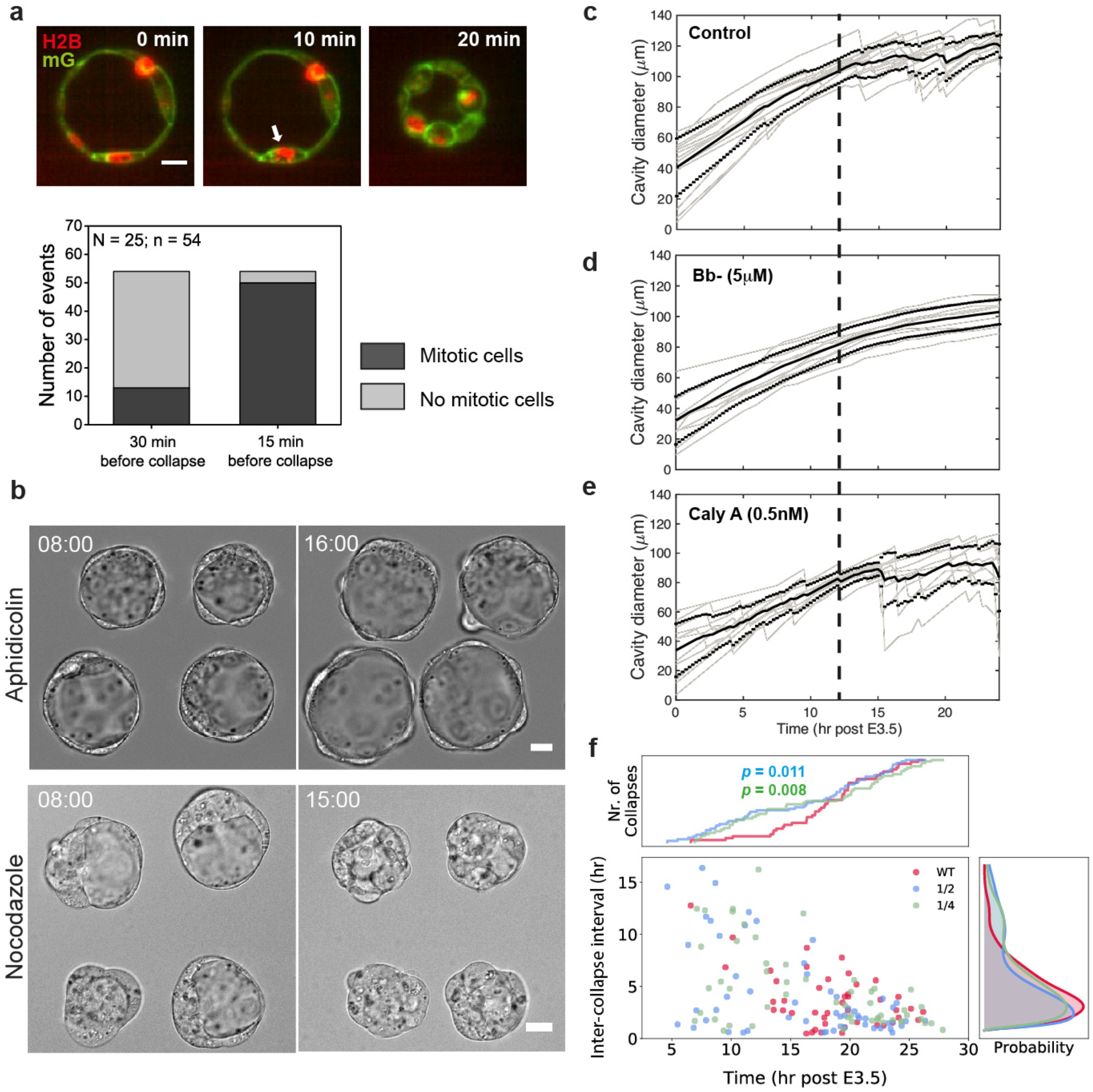
Stretch-induced tissue stress and mitotic cell rounding collectively lead to blastocyst collapse. **a.** Blastocyst collapse is triggered by mitotic cell rounding. Top panel: representative image of an H2B-mCherry/mG expressing blastocyst, showing mitotic cell rounding (white arrow) prior to blastocyst collapse. Bottom panel: quantification of mitotic and non-mitotic events prior to blastocyst collapse, showing that the collapses are highly correlated with preceding mitotic events. Scale bar: 20 μm. **b**. Top panels: representative images of half blastocysts which, when treated with aphidicolin (2 μg/ml), expanded without collapse even after 16 hr post E3.5. Bottom panels: representative images of nocodazole-treated half blastocysts showed constant leakage and overall failed blastocyst expansion. Scale bars: 20 μm. **c-e**. Cavity diameter as a function of time (post-E3.5) for embryos treated with 5 μM of Bb-(*N* = 11) **(d)** and 0.5 nM of Calyculin A (*N* = 11) (e) compared to the control embryos (*N* = 14) **(c)**. Solid black lines denote mean diameter, with ±1SD denoted by dotted lines. Gray lines represent tracks of cavity size evolution for individual embryos. Bold dashed lines are guides to the eye indicating 12 hr post E3.5. **f**. Inter-collapse interval (ICI) plotted as a function of time (post E3.5) for embryos of various sizes. Plot shows more collapse events at early blastocyst stage for the reduced blastocysts compared to the WT (1×) embryos. Kolmogorov-Smirnov test for the number of collapse events compared to the WT embryos: *P* = 0.011 for 1/2 embryos and 0.008 for 1/4 embryos. *N* = 14, 21, 16 for WT, 1/2 and 1/4 embryos, respectively.

In contrast with observations during the late blastocyst stage, we rarely find collapse events in early to mid-blastocyst stage despite extensive cell divisions (Fig. 1a), suggesting that stretch-induced cell stiffening is necessary to ‘prime’ junctional rupture upon mitosis. Consistent with this, our measurements show that the CT of mitotic cells in the early blastocysts is lower (~1400 pN/μm) compared to that in the late blastocysts (~1700 pN/μm, close to those measured in cell culture^18^, Extended Data Fig. 4f). To test this further, we abolished actomyosin contractility using Bb- or enhanced it by calyculin A; the former suppressed collapse while the latter promoted it (Fig. 3c-e). Furthermore, reduced blastocysts, which exhibit higher CT (Fig. 1e), displayed earlier and more frequent collapse compared to the whole blastocysts (Fig. 3f, Extended Data Fig. 5a), supporting the hypothesis.

Together, these findings suggest that blastocyst size is governed autonomously by feedback mechanisms operating across scales, from molecular to cellular and tissue level (Fig. 4a). During blastocyst expansion, the lumenal pressure acts as a global mechanical signal to drive and coordinate cell stretching and generate mechanical stress, which reinforces junctional maturation at the molecular scale through mechanosensing. As the tissue tension reaches a critical threshold, cell-cell adhesion cannot be sustained during mitotic entry, which then triggers junctional rupture and fluid leakage, leading to collapse of the blastocyst cavity. As the pressure is released, the blastocyst heals and the whole process repeats itself so that the steady-state blastocyst size remains constant.

**Figure 4:**
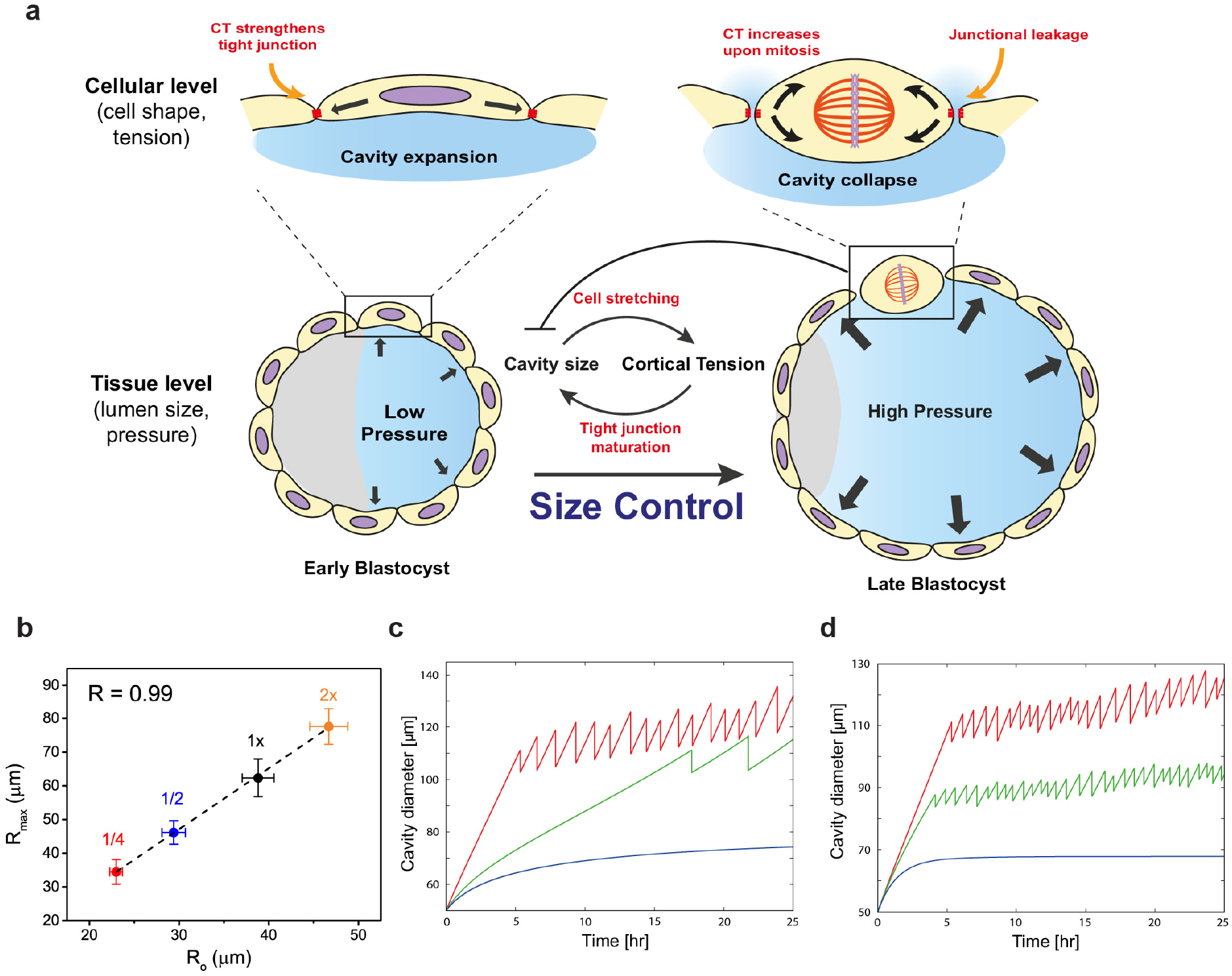
Physical model predicts the change in blastocyst size and oscillation dynamics with perturbation of cortical tension, **a.** Schematic representing how multi-scale feedback interactions between lumenal pressure and tissue mechanics control blastocyst size. Lumen expansion generates mechanical stress on TE cells and triggers mechanosensing and reinforcement of junctional integrity. When the cortical tension reaches a threshold above which cell-cell adhesion cannot be sustained, mitotic entry induces junctional rupture and collapse of the blastocyst cavity, thereby maintaining a steady-state blastocyst size. **b.** Steady-state blastocyst radius as a function of initial blastocyst radius (prior to cavitation), across various reduced systems (red: 1/4 embryos, *N* = 16; blue, 1/2 embryos, *N* = 16; black, 1× embryos, *N* = 22; orange: 2x embryos, *N* = 7). Plot shows a strong correlation (Pearson correlation *R)* between the two parameters, as predicted by the hydraulic model. **c.** Theoretical simulation predicts a delay or abolishment of steady-state oscillations in response to leakage through impaired tight junctions. *J_leak_* = 0 μm/hr (red), *J_leak_* = 5.5 μm/hr (green), = 7.5 μm/hr (blue). **d.** Theoretical simulation further predicts earlier onset and more frequent collapses in response increased CT and tissue stiffness. *E* = 5 kPa + 2.5*σ* and *E_p_* = 50 kPa (red), *E* = 10 kPa + 3*σ* and *E_p_* = 64 kPa (green), and E= 20 kPa + 5σand *E_p_* = 84 kPa (blue).

Our experimental findings are consistent with a recently proposed theory for size control of multi-cellular hollow cysts, using hydraulically-gated oscillations^19^. A key prediction of the theory is that the ultimate size of a hollow blastocyst (*R*_max_) scales linearly with the initial size of the blastocyst (*R*_o_), in the absence of any net increase in tissue volume, i. e. *R*_max_ ~ (1+*σ*_c_/*E*) *R*_o_ where *σ*_c_ is the TE rupture stress, *E* the elastic modulus of the TE epithelium. Using blastocysts of various reduced sizes, we confirmed this linear scaling experimentally (Fig. 4b). Furthermore, using the lumenal pressure measurements and the rupture stress, we estimate *E* ~ 20 kPa, close to those measured for a cellular monolayer^20^. The theory (Supplementary Note) also predicts that leaky cell junctions lead to a delay and a lower frequency of collapse (Fig. 4c), while an increase in the CT (and hence *E)* leads to earlier onset of and more frequent collapses (Fig. 4d), consistent with our observations (Fig. 3c-e).

Additionally, the theory predicts that impaired tight junctions due to reduced actomyosin activity and CT will lead to a reduction in blastocyst size. This would make the behavior of a multi-cellular tissue shell stand in sharp contrast to that of single cells in which actomyosin abolishment leads to cell swelling^21^. We experimentally tested this via pharmacological perturbation of CT, using Bb-or Y27632, a Rho-kinase inhibitor that decreases actomyosin contractility. We found that both these treatments led to reduced cavity expansion rate and smaller size in a dose-dependent manner (Fig. 5a, Extended Data Fig. 6a, c, d). Direct disruption of tight junctions by CPE also resulted in reduced expansion rate and size (Fig. 5b, Extended Data Fig. 6f). Additionally, genetic perturbations in maternal homozygous KO and zygotic heterozygous (*m^−/−^Myh9^+/−^*) embryos, which are known to have reduced CT^22^, displayed a similar reduction in expansion rate and blastocyst size (Fig. 5a, Extended Data Fig. 6a). Furthermore, maternal heterozygous loss of *Myh9* was also sufficient to induce the same effect (Extended Data Fig. 7a-d). In all these perturbations, the total cell number is not significantly different from that in WT embryos (Extended Data Fig. 6b, e, g), suggesting that the change in cavity growth dynamics is not due to reduced cell proliferation that could impact the rate of blastocyst expansion, which approximately scales with the total cell number (Extended Data Fig. 5b).

**Figure 5:**
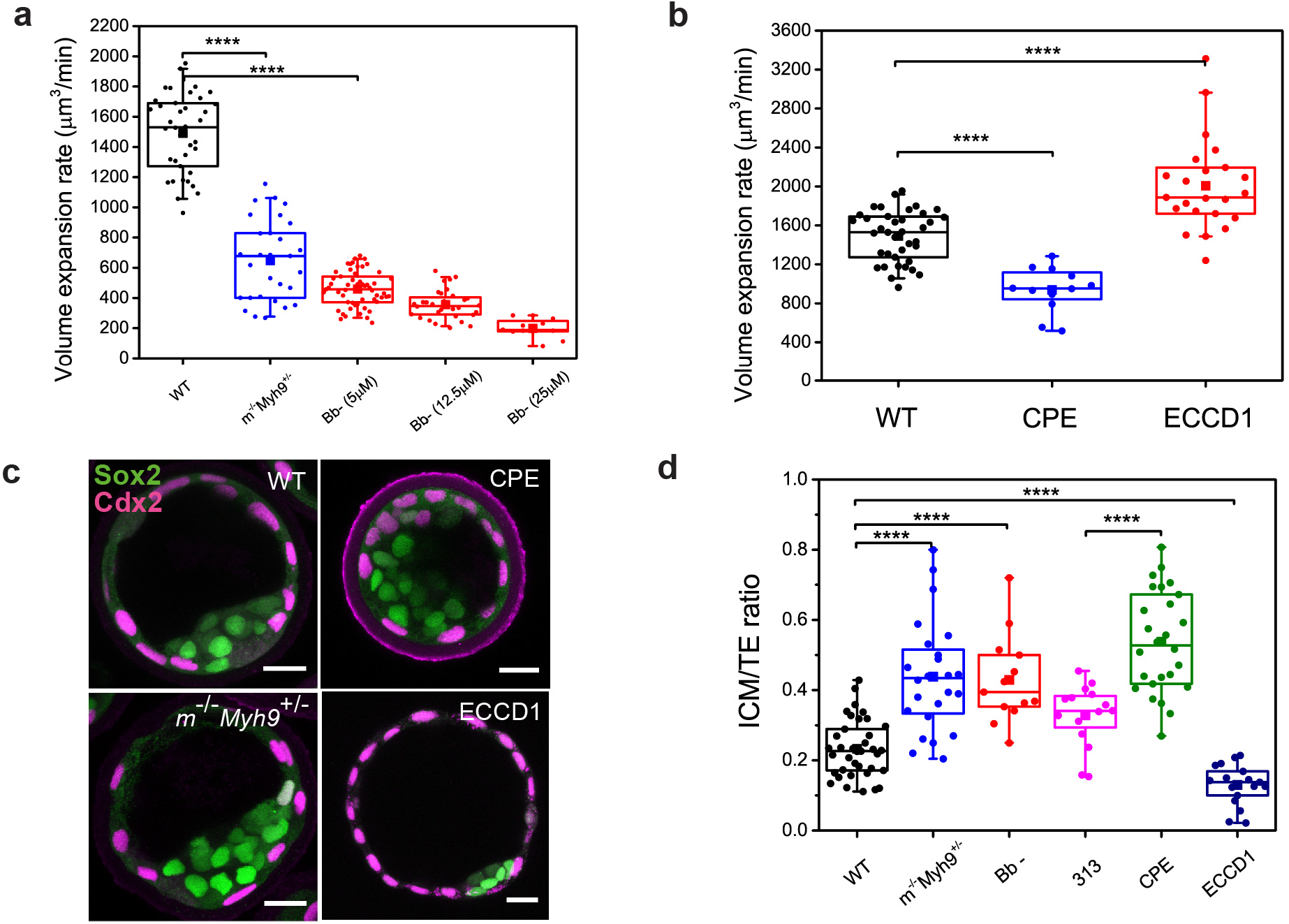
Lumenal pressure couples cell positioning and fate specification. **a.** Volume expansion rate in maternal homozygous KO and zygotic heterozygous *(m^−/−^Myh9^+/−^*) embryos (blue, *N* = 12) and Bb-treated embryos (red, *N* = 15, 8 and 4 for 5, 12.5 and 25 μM, respectively) compared to the control embryos (black, *N* = 23). **b.** Volume expansion rate in CPE-treated (blue, *N* = 12) and ECCD1-treated (red, *N* = 23) embryos compared to the control embryos (black, *N* = 23). **c.** Immunostaining of late blastocyst stage (E4.25) wild-type, *m^−/−^Myh9^+/−^*, CPE and ECCD1-treated embryos showing TE (Cdx2, magenta) and epiblast (Sox2, green) fate. Scale bars: 20 μm. **d.** ICM/TE ratio for E4.25 wild-type (black, *N* = 37), *m^−/−^Myh9^+/−^* (blue, *N* = 24), Bb-treated embryos (red, *N* = 13), CPE-treated embryos (green, *N* = 26) and CPE-control 313, magenta, *N* = 16), and ECCD1-treated embryos (dark blue, *N* = 17).

The theory also predicts that softening (stiffening) the TE shell will lead to an increase (decrease) in the cavity expansion rate and final blastocyst size. Consistent with this, treatment of the blastocyst with ECCD1, E-cadherin-blocking antibody, led to increased cavity growth rate and size (Fig. 5b, Extended Data Fig. 6f), suggesting that weakening of adherens junctions but not tight junctions (Extended Data Fig. 8) could reduce the overall tissue stiffness^23^ that constraints lumenal expansion. Contrariwise, treatment with lysophosphatidic acid (LPA) and calyculin A, both of which activate cortical contractility^24^, led to reduced expansion rate and cavity size (Extended Data Fig. 9), possibly through tissue stiffening. Together, these findings suggest that tissue stiffness might be optimally tuned by cell contractility and adhesion to allow for lumen growth. Collectively, changes in CT, tight junction integrity and tissue stiffness lead to changes in the steady-state blastocyst size, in agreement with the predictions of the model.

Since changes in blastocyst size with unchanged total cell number necessarily impact tissue architecture, we asked if ICM/TE cell allocation would be affected in the perturbed embryos. Remarkably, we found that in embryos with smaller cavities (Bb-, CPE and *Myh9^+/−^*), more cells were allocated to the inner layer of the blastocyst, resulting in a higher ICM to TE ratio (Fig. 5c-d, Extended Data Fig. 7a, e), while those with larger cavities (ECCD1) exhibited a decreased ICM/TE ratio (Fig. 5c-d). While reduced contractility (Bb, *Myh9^+/−^*) may lead to different cell sorting behavior^22^, our data suggest that a change in lumen size due to changes in fluid pressure (CPE) or TE stiffness (ECCD1) can independently lead to altered lineage composition. Of note, *m^+/−^Myh9^+/−^* mutants occasionally failed to spatially segregate into the inner and outer cells (Extended Data Fig. 7f), suggesting that the embryos were developmentally delayed when early Cdx2-positive cells in the ICM had yet to commit to ICM fate^25^. These findings point to the key role of the lumen in defining tissue dimension and pattern. In the mouse pre-implantation embryo, cell position is possibly recognized by apical domain acquisition^26^, that in turn leads to differential Yap signaling and cell fate specification^27^. This experimental outcome is also in line with reports that TE and ICM cell fates remain reversible until the 32-cell and 64-cell stage, respectively^28–30^.

Our study has identified a global mechanical signal, lumenal fluid pressure as a simple regulator that rapidly propagates mechanical information to coordinate cellular response (e.g. change in geometry and CT) and mediate tissue self-organisation (e,g, cell fate) across supra-cellular and sub-cellular scales. Our choice of mouse blastocyst allowed us to probe how feedback signals driven by hydraulic pressure propagate through the system in a minimal setting, as there is no net tissue volume change during this process (cells undergo cleavage). However, such lumen-mediated mechanism may also regulate organ size control at later development, as long as the fluid expansion rate is sufficiently high compared to the tissue growth rate. Given the widespread presence of fluid-filled lumena in epithelial tissue morphogenesis such as in the lung and kidney, it would seem natural to explore hydraulically mediated aspects of tissue size, shape and cellular fate in such systems.

## Acknowledgements

We are grateful to members of the Hiiragi lab and the European Molecular Biology Laboratory (EMBL) animal facility for their support. We thank Mikio Furuse for the plasmids GST-C-CPE/CPE313, and Takafumi Ichikawa and the protein purification facility (EMBL) for protein purification. We thank Dimitri Fabrèges for help with cell volume segmentation, and Esther Jeong Yoon Kim for generating the mouse lines. We thank A. Aulehla, A. Diz-Muñoz, D. Gilmour, F. Graner, E. Hannezo, S. Hopyan, P. Liberali, R. Prevedel and J. Solon for critical reading and constructive feedback on the manuscript. We thank Tomohito Higashi, Ann Miller, and Cornelia Schwayer for kindly sharing with us the protocol for Fluozin leakage assay. C.J.C. and G.M. are supported by fellowships from the EMBL Interdisciplinary Postdocs (EIPOD) under Marie Sklodowska-Curie Actions COFUND (grant number 664726). The Hiiragi laboratory is supported by EMBL, German Research Foundation, and the European Research Council (ERC Advanced Grant “SelforganisingEmbryo”, grant agreement 742732). T.R.-H. was supported by the Simons Foundation. L.M. thanks the MacArthur Foundation and the Radcliffe Institute for support.

## Author Contributions

C.J.C conceived the project and designed the experiments, and wrote the manuscript with input from all authors. C.J.C and M.C. performed and analysed all the experiments. G.M. helped with the ICI quantification. R.J.P. assisted with the initial test runs of the micropressure system. T.R.-H and L.M. created the biophysical model, performed simulations and suggested experimental tests of the model. L.M. contributed to the writing of the manuscript. T.H. supervised the study, helped design the project, performed the oocyte recovery experiments and contributed to the writing of the manuscript.

---

**Extended Data Figure 1:**
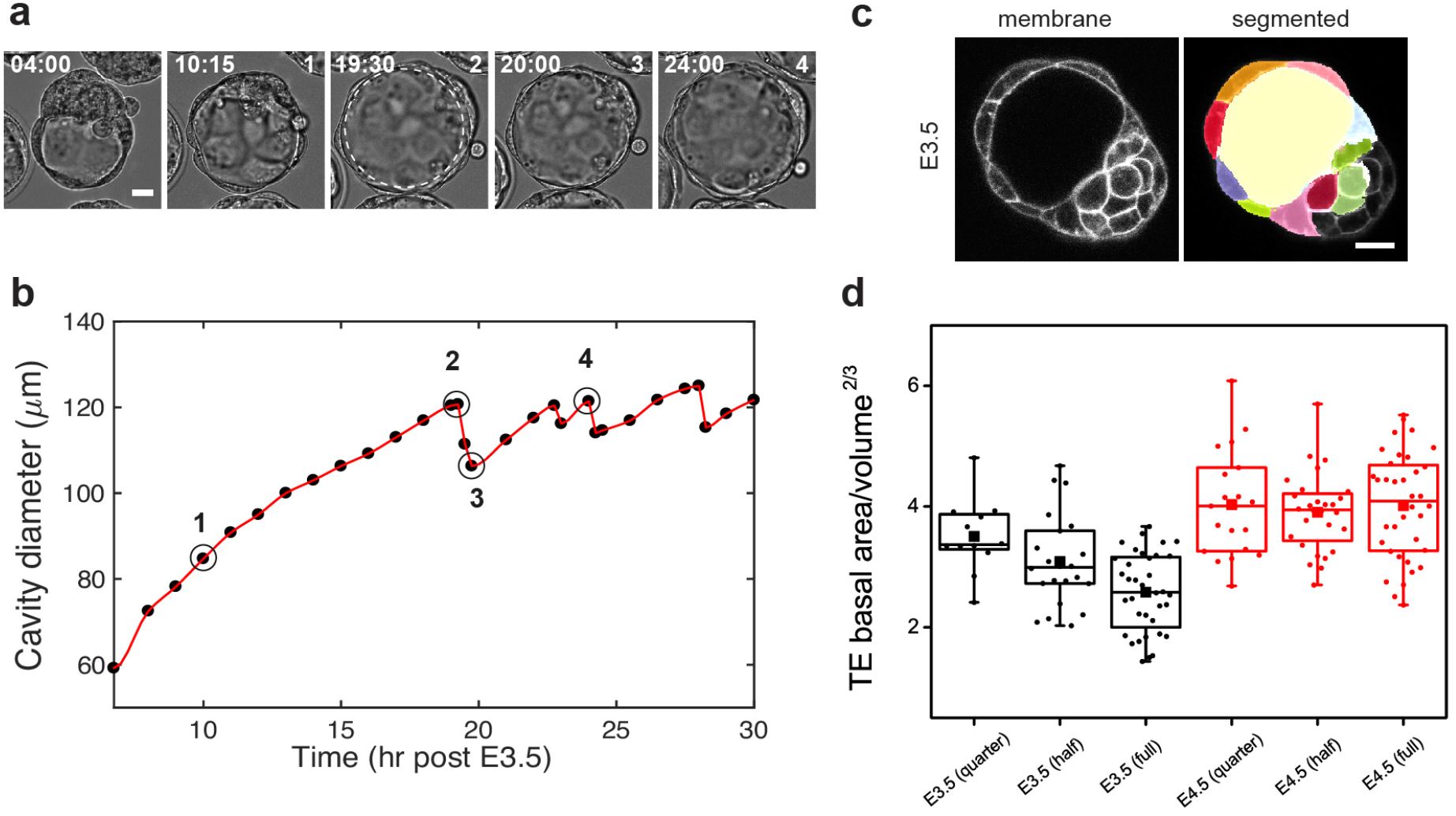
Blastocyst size is controlled by an embryo-autonomous mechanism. **a**. Representative images of ZP-free embryos undergoing blastocyst development. Scale bar: 20 μm. Dotted line denotes the cavity. Time (hh:mm). **b**. Cavity diameter as a function of time (post E3.5) for embryo in **(a)**, showing similar cycles of blastocyst collapse and re-expansion as in ZP-intact embryos. **c.** Representative images of E3.5 blastocyst expressing mTmG (left) and after segmentation (right). Scale bar: 20 μm. **d.** TE basal area normalized to cell volume in various reduced systems at E3.5 (black, *N* = 11, 20 and 20 for 1/4, 1/2 and 1× embryos, respectively) and E4.5 stage (red, *N* = 16, 21 and 22 for 1/4, 1/2 and 1× embryos, respectively).

**Extended Data Figure 2:**
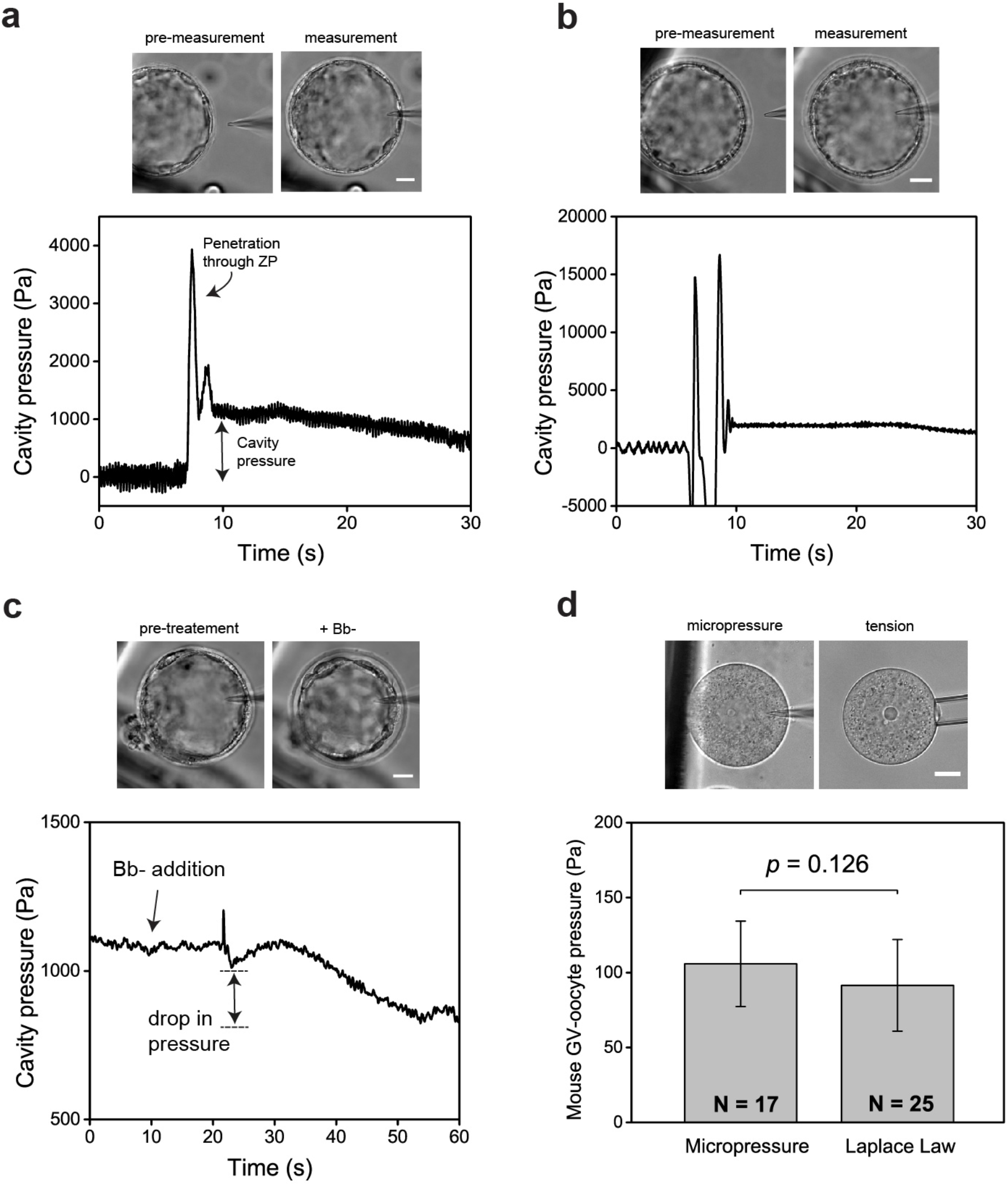
Quantifying cavity pressure in mouse embryos using a micropressure probe. **a-b**. Characteristics of successful cavity pressure measurements. Top panels: Representative images showing mouse blastocysts before and during pressure measurement. The baseline reading is close to 0 when the microelectrode (~0.5 μm) is not in contact with the blastocysts. A transient pressure spike is recorded as the tip of the microelectrode penetrates the ZP. This is followed by a stable reading for 5~10 s before some potential leaks occur, which may cause a gradual drop in the reading with time. **c-d**. Proof-of-principle experiments to verify the pressure measurement: Addition of Bb-(25 μM) leads to transient reduction in cavity diameter and pressure within 10 s. **(c)**. **d**. The accuracy of the device was further demonstrated by comparing the cytoplasmic pressure of mouse oocytes at the Germinal Vesicle (GV) stage, measured directly by the micropressure probe and indirectly by tension measurement and Laplace’s Law. The data obtained by both approaches are not significantly different. *N* = number of oocytes. Scale bars: 20 μm.

**Extended Data Figure 3:**
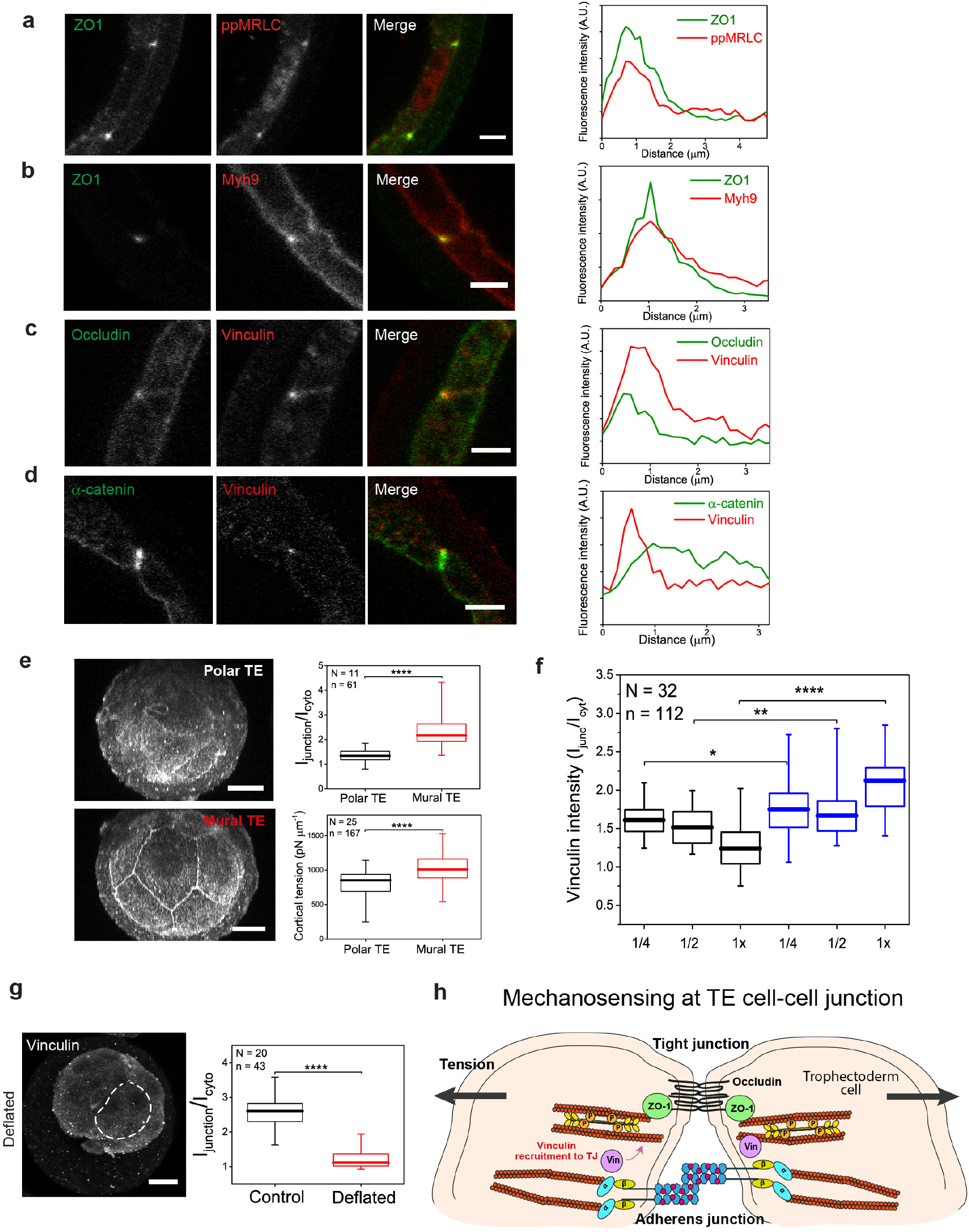
Vinculin are recruited towards the tight junctions in a tension-dependent manner during blastocyst expansion. **a-b**. Immunostaining of wild-type embryos at late blastocyst stage (E4.25) showing co-localization of ZO1 (green) with ppMRLC (red) **(a)** and Myh9 (red) **(b)**. Line scans are plotted on the right. **c-d**. Immunostaining of wild-type embryos at late blastocyst stage (E4.25) showing co-localization of vinculin (red) with occludin (green) **(c)**, and non-localization with α-catenin (green) **(d)**. Line scans are plotted on the right. Scale bars 5 μm. **e**. Box plot (top right) of vinculin intensity ratio (junctional versus cytoplasmic) in polar and mural TE cells in blastocysts at E4.25 stage. Vinculin are more expressed at the mural TE cell-cell junctions compared to those at the polar TE cell-cell junctions (left panels), which correlate with the higher cortical tension measured for the mural TE cells compared to those of polar TE cells (bottom right). Scale bars: 20 μm. **f**. Vinculin intensity ratio (junctional versus cytoplasmic) in various reduced systems, at early (E3.75, black, *n* = 58, *N* = 15) and late (E4.25, blue, n = 54, *N* = 17) blastocyst stage. Vinculin signal intensity is higher in reduced blastocysts compared to the whole embryos at E3.75, and converges to a higher value at E4.25 across all blastocysts, in line with the temporal change in TE cortical tension (Fig. 1e). g. Left: Immunostaining of mechanically-deflated blastocyst (E4.25) showing weak vinculin localization at the TE cellcell junctions following blastocyst deflation. Dotted line denotes the cavity. Scale bar: 20 μm. Right: Box plot of vinculin intensity ratio (junctional versus cytoplasmic) in mural TE cells of control vs. deflated blastocysts. **h**. Schematic showing force-dependent vinculin binding to the tight junctions during blastocyst development as the TE cells get increasingly stretched due to cavity expansion. *n* = number of cells, *N* = number of embryos. \*\*\*\**P*< 0.0001, \*\**P*< 0.01, \**P*< 0.05.

**Extended Data Figure 4:**
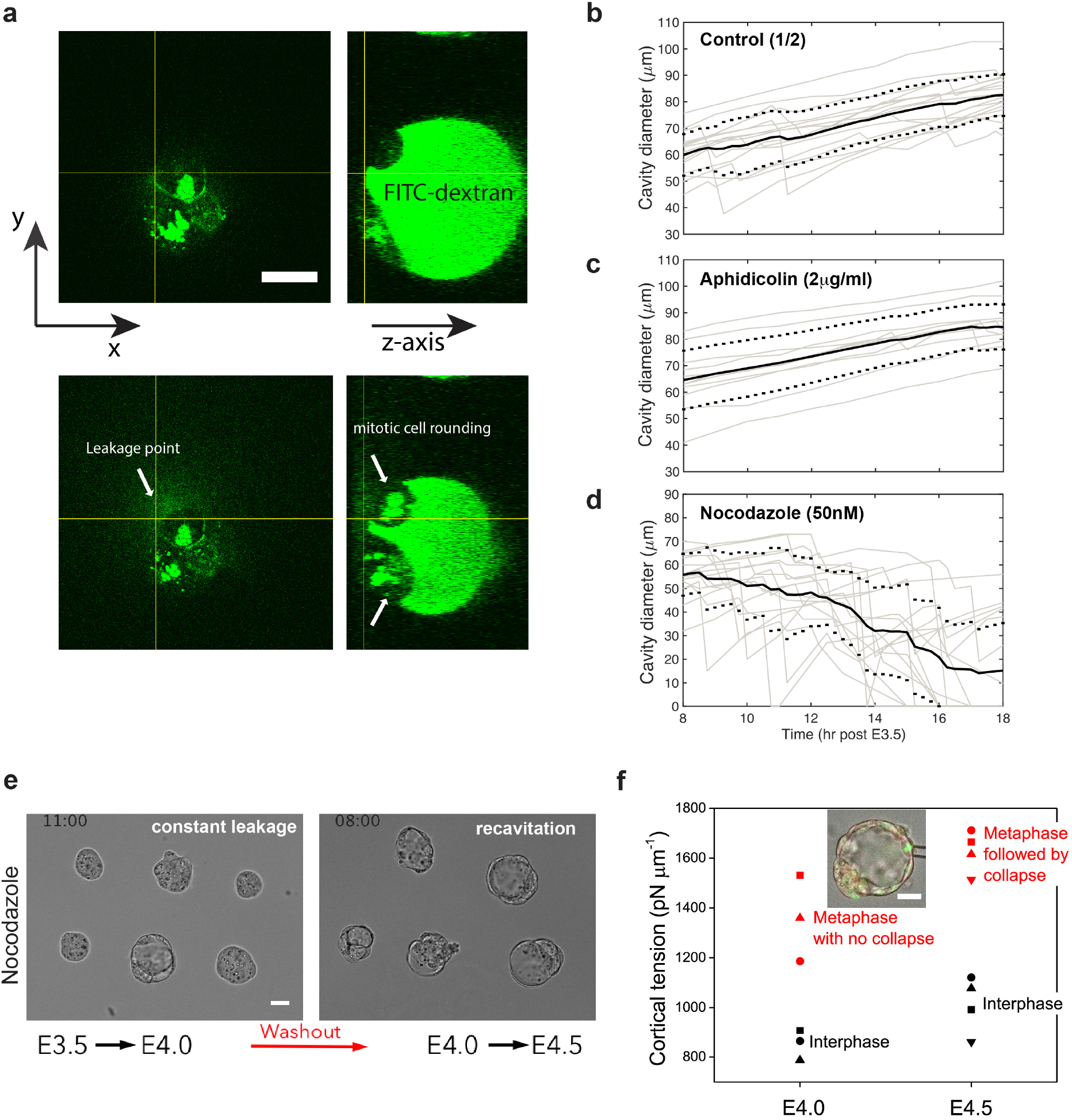
Leakage of blastocoel fluid during TE cell divisions leads to collapse of blastocysts at mature stage. **a**. Representative image of blastocyst (E4.25) showing blastocoel loaded with 4 kDa FITC-dextran dye, prior to (top panels) and after (bottom panels) blastocyst collapse. Dye leakage following blastocyst collapse was observed to occur preferentially at cell-cell junctions between mitotic cells (white arrows). Scale bar: 20 μm. **b-d**. Cavity diameter as a function of time (post E3.5) for 1/2 embryos treated with 2 μg/ml of aphidicolin (*N* = 11) **(c)** and 50 nM of nocodazole (*N* = 17) **(d)** compared to the control 1/2 embryos (*N* = 17) **(b). e**. Representative images showing that 1/2 blastocysts that failed to re-expand upon initial collapse when treated with 5 μM of nocodazole, can recavitate upon washout. Scale bar: 30 μm. **f**. Cortical tension of TE cells measured during interphase (black) and metaphase (red) at mid- (E4.0) and late (E4.5) blastocyst stage. Different symbols correspond to different cells being measured. Inset: Representative image showing tension measurement for a mitotic TE cell of a blastocyst at E4.5. Scale bar: 30 μm.

**Extended Data Figure 5:**
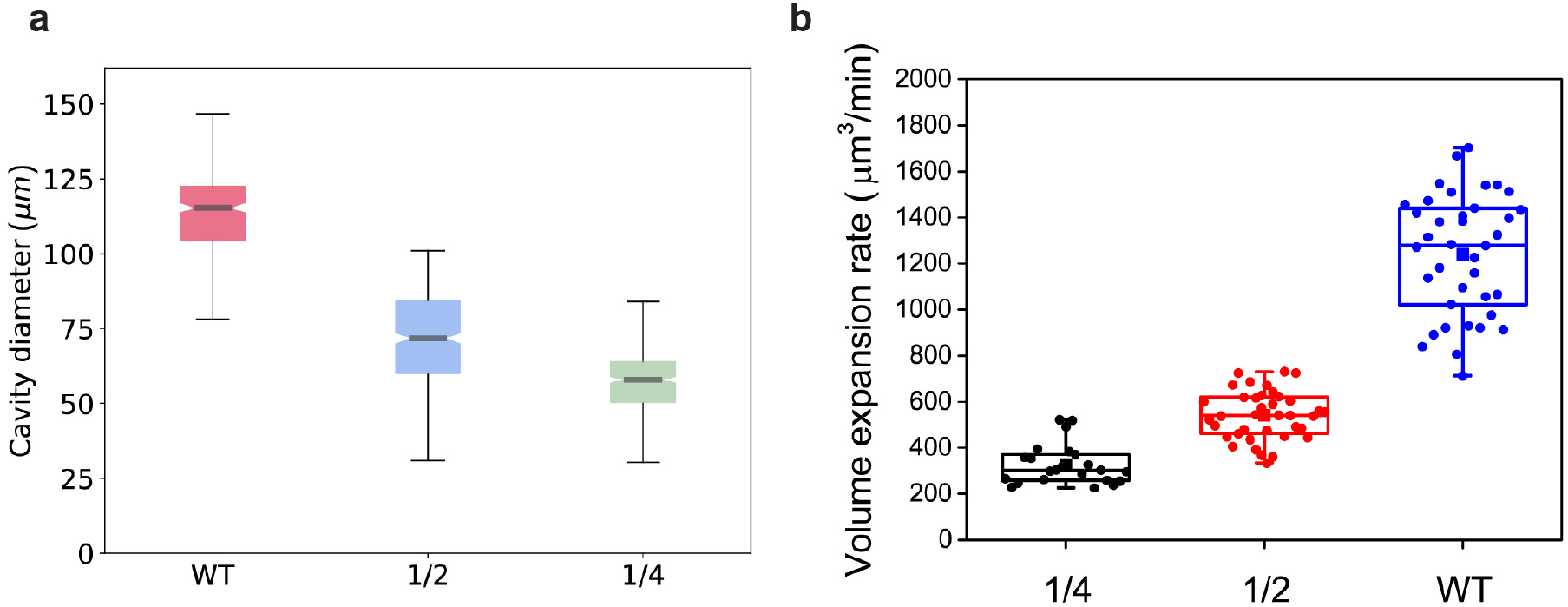
Cavity growth dynamics of reduced versus full blastocysts. **a**. Cavity diameter at the plateau stage for full blastocysts (without ZP) and reduced blastocysts (1/2, blue; 1/4, green). **b**. Plot of volume expansion rate for reduced and full blastocysts shows that the blastocyst expands at a rate that scales approximately with the initial number of cells. *N* = 14, 21, 16 for 1×, 1/2 and 1/4 embryos, respectively.

**Extended Data Figure 6:**
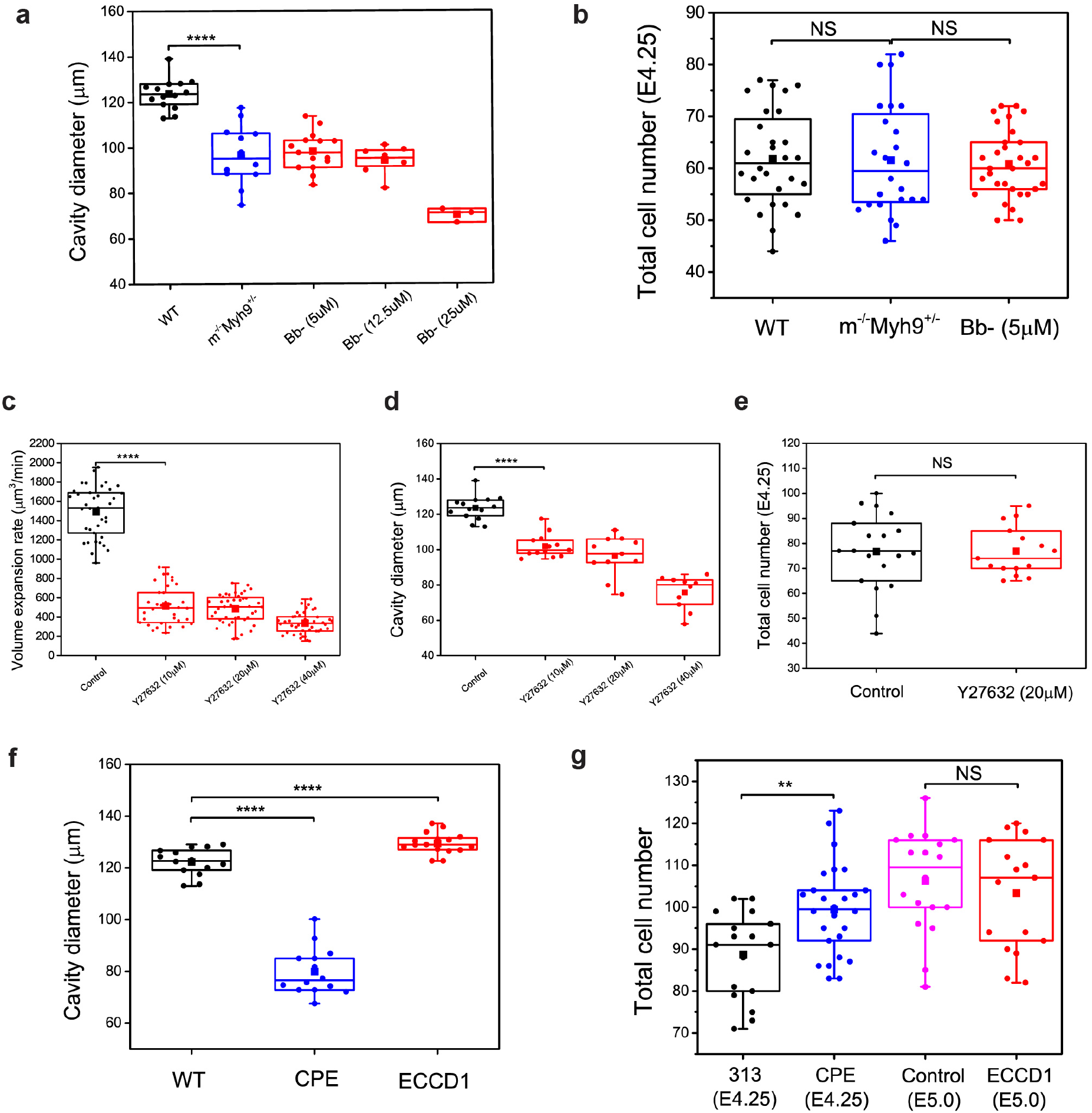
Control of blastocyst size by cortical tension and tight junction permeability. **a**. Maximum cavity diameter reached during time period E3.5 to E4.5 for wild-type (black, *N* = 14), *m^−/−^Myh9^+/−^* (blue, *N* = 12), and embryos treated with Bb-(red) in various concentrations (5 μM, *N* = 15; 12.5 μM, *N* = 8; 25 μM, *N* = 4). These embryos have similar cell number **(b)**, showing that the change in cavity size is not due to reduced cell proliferation rate. *N* = 28, 24 and 31 for WT, *m^−/−^Myh9^+/^*, and 5 μM Bb-treated embryos, respectively. **c-e**. Embryos treated with Y-27632, a Rho-kinase inhibitor targeting actomyosin contractility, showed similar reduction in volume expansion rate (c) and cavity size **(d)**, despite the similar cell number compared to the controls **(e)**. *N* = 14, 13, 11 and 10 for controls, 10, 20 and 40 μM Y-27632-treated embryos, respectively in **(c)** and **(d)**. *N* = 19 and 15 for controls and 20 μM Y-27632-treated embryos, respectively in **(e)**. **f.** Maximum cavity diameter reached during time period E3.5 to E4.5 for CPE (blue, *N* = 14) and ECCD1-treated (red, *N* = 16) compared to WT (black, *N* = 14). g. ECCD1-treated embryos have similar cell number compared to their controls, while CPE-treated embryos have a slightly higher cell number, despite having reduced cavity size. *N* = 17, 26, 18 and 17 for 313 (CPE-control), CPE, WT and ECCD1-treated embryos, respectively. \*\*\*\**P* < 0.0001, \*\**P*< 0.01, *NS* = non-significant.

**Extended Data Figure 7:**
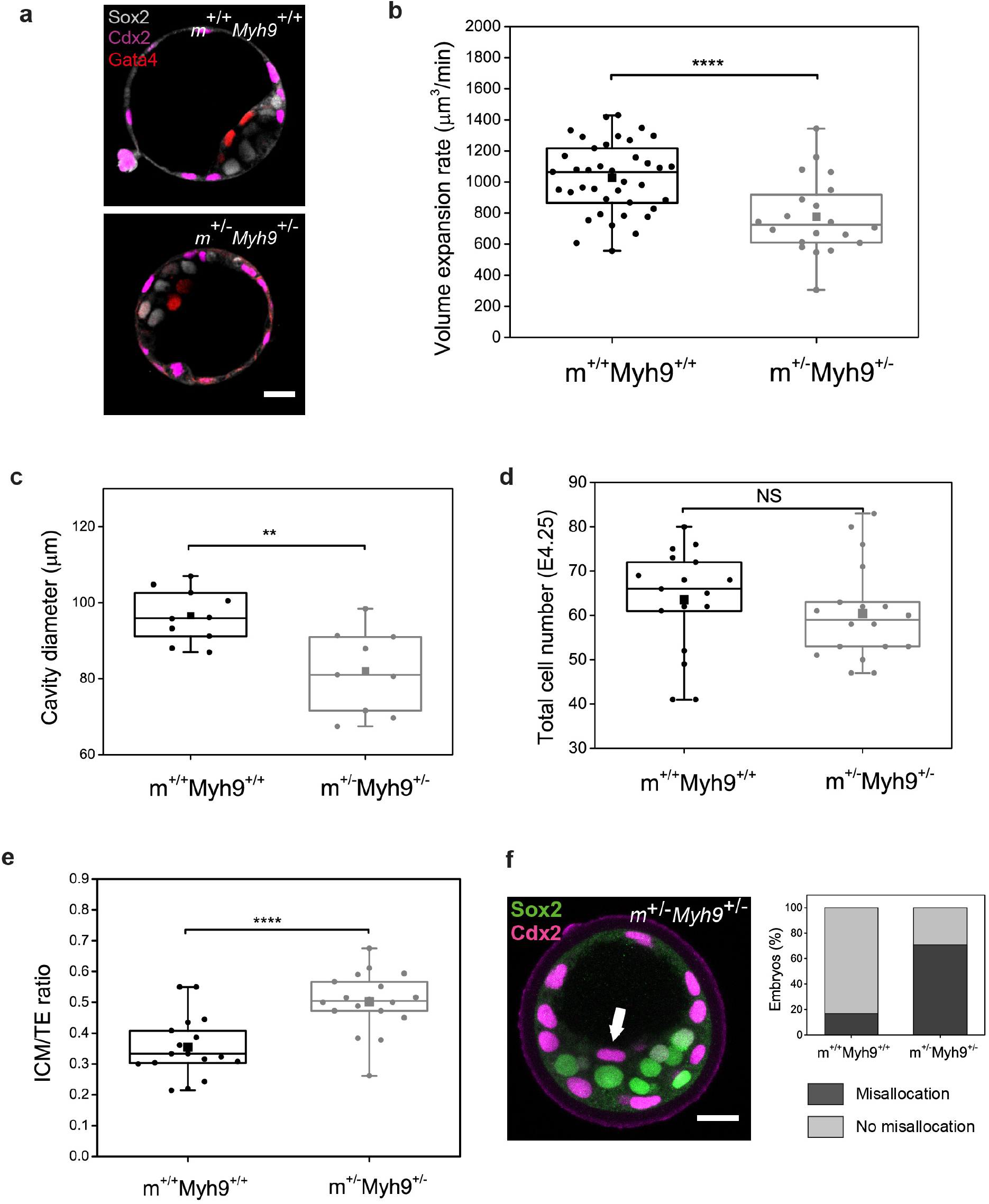
Maternal heterozygous KO and zygotic heterozygous (*m^+/−^Myh9^+/^*) embryos exhibit smaller cavity and increased inner cell mass compared to WT in the same litter-mate. **a**. Immunostaining of late blastocyst stage (E4.25) *m^+/−^Myh9^+/−^* and wild-type embryos in the same litter-mate (*m^+/+^Myh9^+/+^*), showing TE (Cdx2, magenta), epiblast (Sox2, grey) and primitive endoderm (Gata4, red) fate. Scale bar: 20 μm. **b-e**. Volume expansion rate for *m^+/−^Myh9^+/−^* is significantly lower than that of the wild-type *m^+/+^Myh9^+/+^* **(b)**, leading to a smaller cavity size **(c)** and higher ICM/TE ratio (**e**), despite a similar range of cell number in the two cases **(d). (b, c)**. *N* = 9 and 10 for *m^+/+^Myh9^+/+^* and *m^+/−^Myh9^+/−^*, respectively. **(d, e)**. *N* = 17 and 18 for *m^+/+^Myh9^+/+^* and *m^+/−^Myh9^+/−^*, respectively. **f**. Immunostaining of *m^+/−^Myh9^+/−^* mutants (E4.25) showing misallocation of Cdx2-expressing cells in the inner cell mass (white arrow). Scale bar: 20 μm. Right panel: quantification of lineage misallocation in *m^+/−^Myh9^+/−^* mutants. *N* = 35 and 24 for *m^+/+^Myh9^+/+^* and *m^+/−^Myh9^+/−^*, respectively. \*\*\*\**P*< 0.0001, \*\**P*< 0.01, *NS* = non-significant.

**Extended Data Figure 8:**
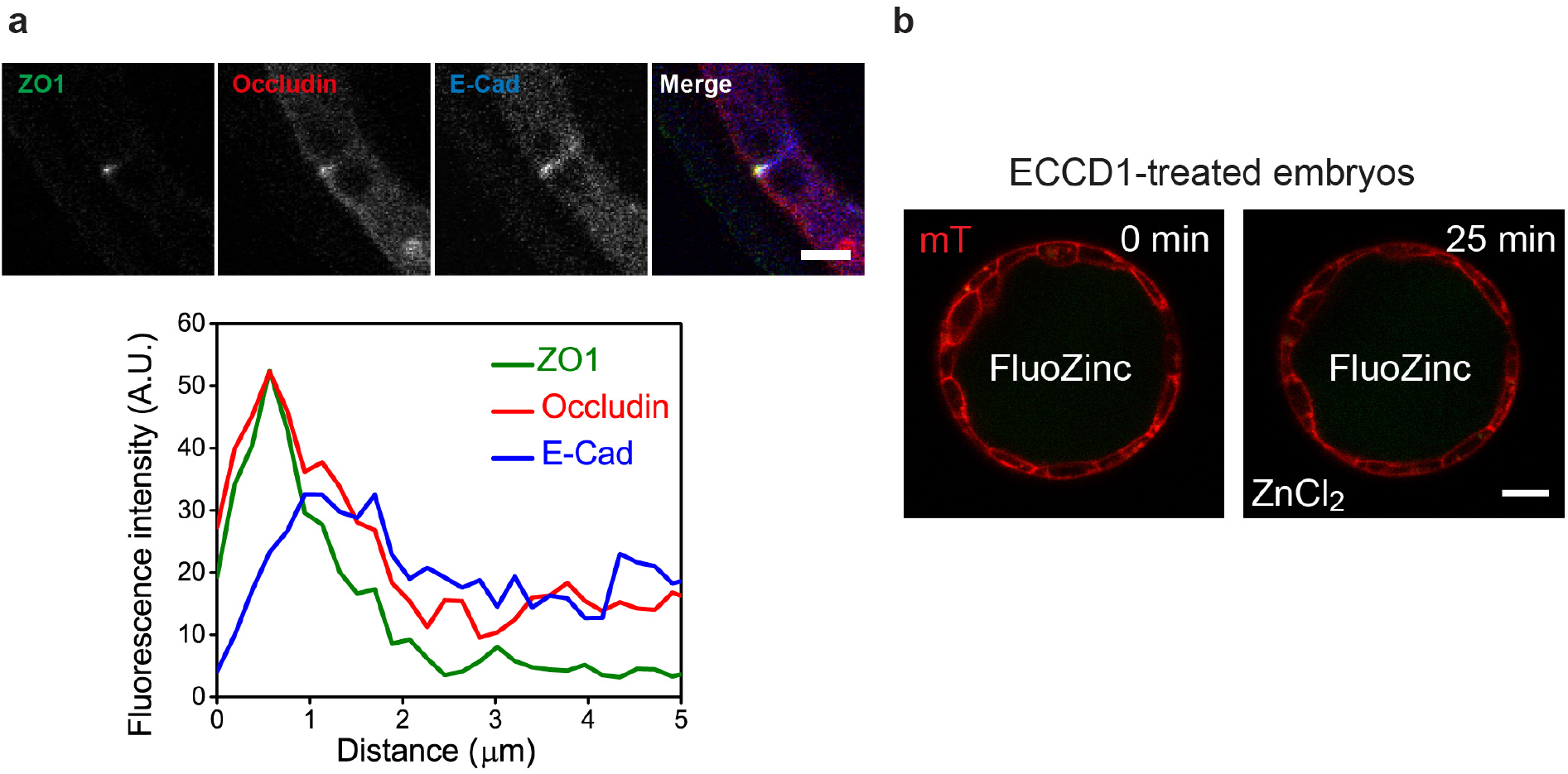
Embryos treated with ECCD1 display intact tight junctions. **a**. Immunostaining of embryos at late blastocyst stage (E5.0), after being treated with ECCD1, an E-Cad blocking antibody. Line scan plot is shown below, with 0 μm indicating the apical-most part of the tight junction. A.U., arbitrary units. Tight junctions appear intact as shown by the decoupling of signals of ZO1 and occludin from E-Cad. Scale bar: 5 μm. **b**. Tight junction leakage assay using Fluozinc dye (100 μM). ECCD1-treated embryos loaded with Fluozinc showed no increased signals within the blastocoel with time when immersed in ZnCl_2_ (200 μM), suggesting that the tight junctions are properly sealed and the functional integrity of tight junctions are unaffected by impaired adherens junctions. Scale bar: 20 μm.

**Extended Data Figure 9:**
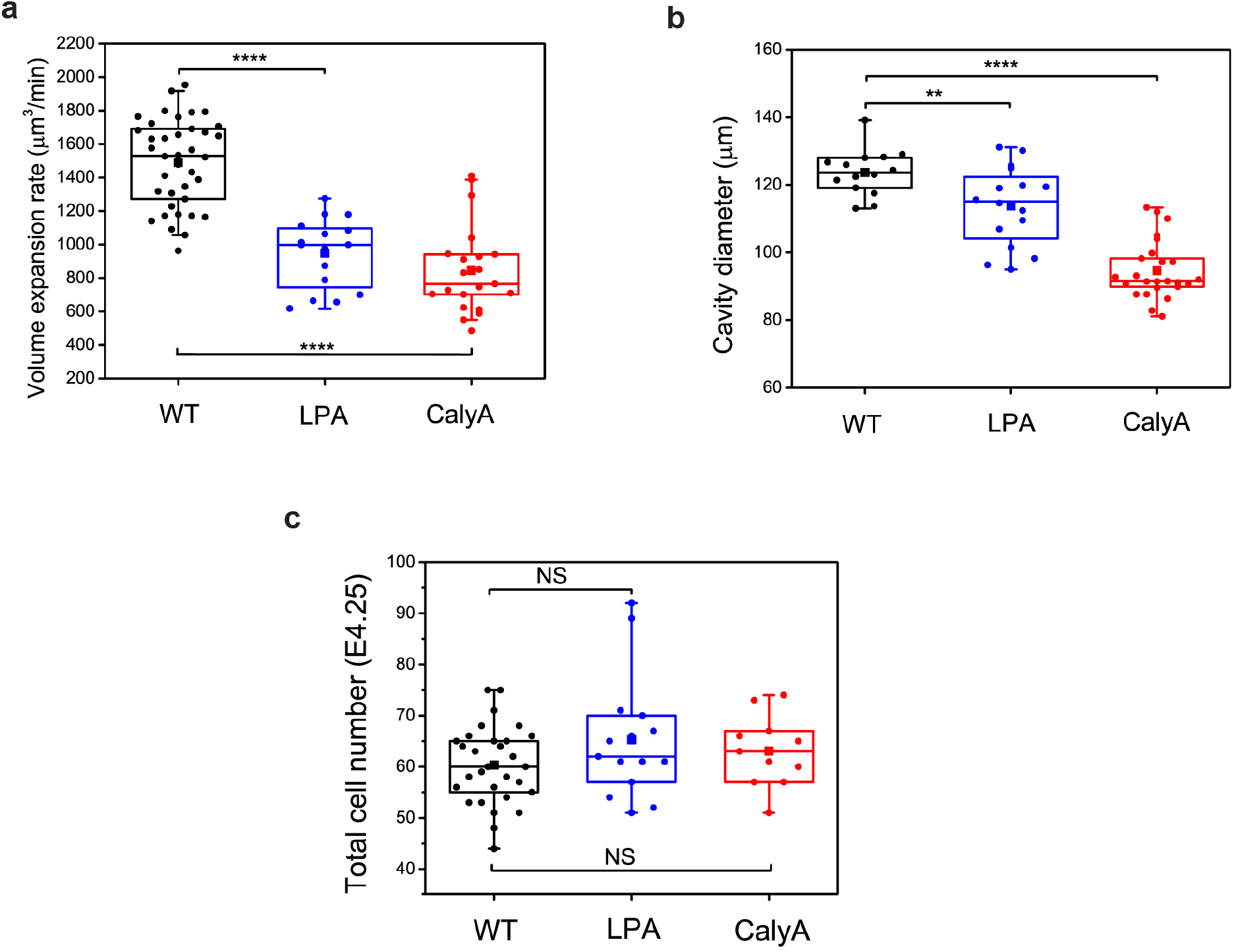
Increased cortical tension leads to reduced blastocyst expansion rate and size. **a**. Volume expansion rate in embryos treated with 20 μM lysophosphatidic acid (LPA, blue, *N =* 16), and 0.5 nM calyculin A (CalyA, red, *N =* 21) compared to the control embryos (black, *N* = 38). Both LPA and CalyA are known to activate cortical contractility. **b**. Maximum cavity diameter reached for WT *(N* = 14), LPA *(N* = 16) and CalyA *(N* = 25). **c**. Total cell number for control embryos *(N* = 30) and embryos treated with LPA *(N* = 15) and CalyA *(N* = 11) is similar, indicating that the change in expansion rate and cavity size is not due to reduced cell proliferation rate. \*\*\*\**P*< 0.0001, \*\**P*< 0.01, *NS* = non-significant.

## METHODS

### Embryo work

#### Recovery and culture

Animal work was performed in the animal facility at the European Molecular Biology Laboratory, with permission from the institutional veterinarian overseeing the operation (ARC number TH11 00 11). The animal facilities are operated according to international animal welfare rules (Federation for Laboratory Animal Science Associations guidelines and recommendations). Mice were used from 8 weeks old onwards.

Embryos were recovered from superovulated female mice mated with male mice. Superovulation was induced by intraperitoneal injection of 5 international units (IU) of pregnant mare’s serum gonadotropin (PMSG; Intervet Intergonan), followed by intraperitoneal injection of 5 IU human chorionic gonadotropin (hCG; Intervet Ovogest 1500) 44-48 h later. Embryos were recovered both at the 2-cell-stage (E1.5) or at the 8-cell-stage (E2.5) by flushing oviducts from plugged females with 37 °C KSOMaa with Hepes (Zenith Biotech, ZEHP-050, 50mL) using a custom-made syringe (Acufirm, 1400 LL 23). Embryos were handled using an aspirator tube (Sigma, A5177-5EA) equipped with glass pipettes pulled from glass micropipettes (Blaubrand intraMark). For *in vitro* culture, embryos were transferred to 10 μl droplets of KSOM (Millipore, MR-121-D) in a tissue culture dish (Falcon, 353001) covered with mineral oil (Sigma, M8410 or Acros Organics), and cultured in an incubator with a humidified atmosphere supplemented with 5% CO_2_ at 37 °C.

Embryos were dissected out of *zona pellucidae* (ZP) at 4-cell stage using a holding pipette and a glass needle^31^, or dissolved using pronase (0.5% protease (Sigma, P-8811) in KSOMaa with Hepes and 0.5% PVP-40). ZP-removed embryos were cultured in Petri dishes (Falcon, 351008). For pressure measurement and micropipette aspiration, ZP-removed embryos were transferred to KSOMaa with Hepes covered with oil, in 50 mm glass bottom dishes (MatTek). For time-lapse imaging, embryos were transferred to KSOMaa with Hepes in 0.16-0.19 mm glass bottom dishes (MatTek).

#### Reduced systems

ZP-removed 4-cell-stage embryos were washed in Ca^2+^-free KSOM and aspirated multiple times through a narrow glass pipette (with radius comparable to the whole embryo). Dissociated blastomeres (1/4 embryos) were then washed multiple times in KSOM before being transferred to KSOM for culture. 1/2 embryos were made by placing 2 blastomeres in micro-indented wells filled with KSOM.

#### Mouse lines and genotyping

embryos derived from mating between (C57BL/6xC3H) F1 hybrid strain were used for wild type (WT) embryos. mTmG mice (Gt(ROSA)26Sortm4(ACTB-tdTomato,-EGFP)Luo)^32^ or mG mice (after Cre-mediated excision of mT) were used to visualize plasma membranes, while H2B–GFP or H2B-mCherry mice^33^ were used to visualize nuclei. Genes were deleted maternally using Zp3-cre (Tg(Zp3-cre)93Knw) mice^34^: to generate m^−/−^Myh9^+/−^ embryos, B6C3F1 males were crossed with Myh9^fl/fl^ :ZP3Cre+^/-^ females^22^. To generate m+^/-^Myh9+^/-^ embryos, B6C3F1 males were crossed with Myh9^+/−^ females. To discriminate between WT and mutant embryos in the same littermate, embryos at E4.25 blastocyst stage were genotyped following immunofluorescence staining and imaging. For genotyping of blastocysts, fixed or live blastocysts were lysed, followed by PCR with the following primers: Myh9EKFw3 (5’-TTGCAGCCCTTCTTGACCTA-3’), Myh9EKRv3 (5’-GCCACATCTCAGCCTAGGAT-3’), Myh9EKDel 1 (5’-CGGGAAGGAAGGAGACACTT-3’).

#### Chemical reagents

To reduce contractility, blebbistatin(+), an inactive enantiomere of the inhibitor, or (-), the selective inhibitor of myosin II ATPase activity (Tocris, 1853 and 1852), were diluted to 5, 12.5 or 25 μM in KSOM from DMSO stocks. Y-27632 (Sigma) in H_2_O stocks were diluted to 10, 20 and 40 μM in KSOM. To induce higher cortical tension, calyculin A (Sigma) in DMSO stocks were diluted to 0.5 nM in KSOM, while lysophosphatidic acid (LPA, Sigma) was used at 20 μM. To perturb mitosis, 2 μg/ml of aphidicolin was used to arrest embryos in S-phase, while 50 nM of nocodazole was used to arrest embryos in metaphase. To reduce cell-cell adhesion, ECCD1, an E-Cad blocking antibody was used at 50 μg/ml and added 3 hr prior to E3.5. To induce a smaller cavity, ouabain (Sigma), an inhibitor of Na/K^+^ ATPase that establish osmotic gradient for blastocysts expansion, was used at 500 μM. Tight junction inhibition was achieved using recombinant GST-C-CPE fusion protein, an inhibitor of Claudin4 and 6. GST-C-CPE and its inactive form, GST-C-CPE313, were diluted to 100 μM before being applied to embryos at E3.25. Embryos were incubated with drugs from E3.5 to E4.5, unless otherwise stated.

#### GST-C-CPE and GST-C-CPE313 purification

Plasmids encoding the C-terminal half of *Clostridium perfringens* enterotoxin (C-CPE, aa 184-319) and its deletion mutant (C-CPE313, aa 184-313) were kindly provided by Dr. Mikio Furuse. GST-tagged C-CPE and C-CPE313 were expressed in *E.coli* BL21 and purified as described previously^13^, with slight modifications. *E.coli* cells were sonicated in lysis buffer (50mM Tris, 300 mM NaCl, 5% glycerol, 1 mM DTT, 0.1% Triton X-100, protease inhibitor cocktail, DNase, 5mM MgCl_2_). The proteins were purified by GST-beads (Protino Agarose Glutathione 4B, MACHEREY-NAGEL GmbH), followed by dialysis against PBS by using Slide-A-Lyzer™ Dialysis Cassettes, 3.5K molecular weight cutoff (Thermo Fisher).

#### Immunofluorescence staining

Embryos were fixed using 1% PFA in PBS for 15 min at room temperature, washed with PBS added with Tween20 (Sigma, P7949) (PBST) and 3% BSA (Sigma, 9647), and permeabilized in 0.5% Triton X-100 (Sigma, T8787) in PBS. After washing embryos were left in blocking solution (PBST with 5% BSA) for at least 4 h at 4°C and incubated overnight at 4°C with primary antibody diluted in washing buffer. After washing, embryos were incubated for one hour at room temperature in secondary antibody diluted in washing buffer. Embryos were then washed in washing buffer before imaging.

Primary antibodies against Cdx2 (Sigma, 71-1500) were used at 1:50. Primary antibodies against ZO1 (Invitrogen, 339100), α-catenin (Abcam, ab51032), vinculin (Sigma, V9264), diphosphorylated form (Thr18/Ser19) of the myosin regulatory light chain (Cell Signaling, 3674), myosin heavy chain Myh9 (Cell Signaling, 3403), and Gata-4 (Santa Cruz Biotechnology, sc-9053) were used at 1:100. Primary antibody against occludin (Invitrogen, SA133520187) was used at 1:125. Primary antibody against Sox2 (R&D Systems, AF2018), Cdx2 (Biogenex, MU392A-UC) was used at 1:200.

Secondary antibody targeting mouse Ig coupled to Alexa Fluor 488 (Life Technologies, A21202), Alexa Fluor 555 (Life Technologies, A31570), rabbit Ig coupled Alexa Fluor 546 (Invitrogen, A10040), rabbit Ig Alexa Fluor 488 (Life Technologies, A11008), goat Ig coupled to Alexa Fluor 488 (Life Technologies, A11055), goat Ig Alexa Fluor 555 (Life Technologies, A21432) and Cy5 targeting mouse Ig (Jackson ImmunoResearch, 715-175-150) were used at 1:200. DAPI (Invitrogen, D3751) was used at 1:2000. Rhodamine-coupled phalloidin (Invitrogen, 1810954) was used at 1:500 while Alexa 633-coupled phalloidin (Invitrogen, R415) was used at 1:50.

#### Tension measurement

The method has been described in previous studies^11^. In brief, a microforged micropipette coupled to a microfluidic pump (Fluigent, MFCS) was used to measure the cortical tension of TE cells. To measure TE cortical tension at 32-to 96-cell-stage (E3.5 to E4.5), micropipettes with radii 4-6 μm were used to apply step-wise increasing pressures on these cells until a deformation with the same radius of the micropipette (*R*_p_) is reached. Care is taken to avoid aspiring cell cortex next to the cell nuclei since cell nuclei are known to be stiff^35^ and may introduce artefacts in the measured cortical tension. At steady state, the cortical tension *γ* of the blastomere is calculated based on Young–Laplace’s law: *γ* = P_c_/2(1/R_p_ – 1/R_c_), where P_c_ is the pressure used to deform the cell of radius R_c_

#### Pressure measurement

The 900A micropressure system (World Precision Instruments) was used to make direct measurements of blastocoelic pressure according to the manufacturer’s instructions and as described in previous work^10^. Briefly, a 0.5 or 1-μm micropipette (World Precision Instruments) was filled with 1 M KCl solution, placed in a microelectrode holder half-cell (World Precision Instruments), and connected to a pressure source regulated by the 900A system. The microelectrode was calibrated using a calibration chamber (World Precision Instruments), before mounting it onto a micromanipulator (Narishige) within an inverted Zeiss Axio Observer microscope. Blastocysts were first seeded onto dishes glued with coverslips (Thermo Scientific Menzel, 1130486) to provide a wedge against embryo movement during pressure measurement. The microelectrode was then inserted into the blastocyst and maintained in place for ~10 s before removal. Pressure measurement was calculated as the mean pressure reading during this time interval, with a resolution of 13 Pa. The accuracy of the measured pressure from the micropressure system was validated by cross-checking with the cytoplasmic pressure of mouse oocytes (GV stage), which was calculated using the Laplace’s Law: *P_c_ = 2γ/R* where *γ* is the cortical tension of the oocyte measured by micropipette aspiration, and *R* is the radius of the oocyte.

#### Microscopy

Tension measurements were performed on an inverted Zeiss Axio Observer microscope equipped with a dry **x**40/0.75 Plan-Neofluar objective. For long-term time-lapse imaging of blastocyst development (E3.5 – E4.5), a dry **x**20/0.8 PL Apo DICII objective was used. Images are captured by a Zeiss AxioCam MRm using Zeiss Axiovision software. The microscope is equipped with an incubation chamber to keep the sample at 37°C. Confocal imaging of both fixed and live samples was performed using Zeiss LSM 780 Confocal Inverted Microscope with LD C-Apochromat 40x/1.1 W Corr objective, using Zen 2012 LSM software. Laser lines used are diode 405 nm, argon multi-line 458/488/514 nm, HeNe 561 nm and 633nm. Laser power and digital gain settings were unchanged within a given session to permit direct comparison of expression levels among embryos stained in the same batch. Image stacks were acquired with axial step size of 1 μm or less. Sequential image acquisitions were used to avoid bleed-through artefacts across different channels.

#### Tight junction permeability assay

Embryos were first incubated in KSOM added with 100 μM Fluozinc, 10 μM Ca-EDTA for 8 hours at 32-cell stage to allow the dye to be loaded into the blastocoel by osmotic gradient. The embryos were then washed out, transferred to KSOM and imaged to acquire the fluorescence intensity in the blastocoel. Additional KSOM with 200 μM ZnCl_2_, either with Bb+ (control) or Bb-, was then added to the medium (*t* = 0 min). Tight junction permeability was then assessed by quantifying the increase in fluorescence intensity within the blastocoel with time.

To trace leakage sites in blastocysts undergoing collapse, embryos were first incubated in KSOM added with 4 kDa fluorescein isothiocyanate (FITC)–dextran for 8 hours at 32-cell stage. The embryos were then washed in KSOM at E4.5 blastocyst stage and imaged every 10 min. Upon seeing any mitotic cells in the blastocysts (visible metaphase plate in H2B-GFP expressing embryos), image acquisition was switched to fast mode to trace dye leakage points across the entire blastocyst.

### Data analysis and Statistics

#### Cell shape analysis

Using FIJI, a circle was manually fit onto the cell-medium interface to extract the radius of curvature of the cell *R*_c_. The diameter of the micropipette *R*_p_ was quantified by drawing a line perpendicular to the micropipette tip followed by the use of linescan function. To quantify basal area and cell volume of TE cells during blastocyst expansion, embryos expressing mTmG were imaged with less than 1 μm spacing of confocal slice. TE cells near the equatorial plane of the blastocysts were picked for manual quantification of TE basal area. Following segmentation using a pathfinder algorithm (Dijkstra’s algorithm), cell volume was calculated using the 3D ImageJ Suite plugin in FIJI.

#### Time-lapse analysis

A circle was manually fitted to the blastocyst cavity at the equatorial plane to extract the cavity diameter. Cavity volume was calculated assuming spherical symmetry of the blastocyst cavity, which was confirmed by direct imaging of cavity shape using 4 kDa dextran-FITC dye. The steady-state cavity diameter was extracted from the maximum size reached during blastocyst development from E3.5 to E4.5. Volume expansion rate was calculated from the change in volume per hour in the last few hours before E4.5, excluding time points during the collapse phase. MATLAB (The MathWorks, Natick, MA) was used to plot the temporal evolution of cavity diameter during blastocyst development.

#### ICI quantification

To detect the blastocyst collapses, interpolating splines were fitted to the measured cavity diameter traces. Zero crossings of the 1^st^ derivative then indicate switches from expansion to shrinkage. To exclude spurious diameter drops, the local minima of the 1^st^ derivative were thresholded with 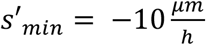. For every trace this method gives a series of collapse event times: *T_collapses_* = {*T*_1_, *T*_2_,…, *T_N_*]. The empirical inter-collapse-interval (ICI) distributions are then simply given by the respective time differences: Δ*T_i_* = *T*_*i*+1_ – *T_i_*. The continuous ICI densities shown in Fig. 1c (right panel) were obtained by Gaussian kernel density estimation of the log-transformed ICIs, the kernel-bandwidth was selected using Silvermans criteria^36^. The whole analysis was done with custom written Python scripts, using tools from the scientific package SciPy.

#### Intensity ratio measurements

Using FIJI, a free-hand line with thickness ~1 μm was drawn along the junctions between two mural TE cells. After measuring the mean gray value for the intensity of the signal at the junctions, the same line was shifted to the cytoplasm of one of the two cells to measure the mean gray value for the cytoplasmic signal. The mean gray value for junctional signal was then normalized by the mean gray value for cytoplasmic signal.

#### Line scan and skeletal analysis

Using FIJI, we picked confocal slices near the equatorial plane of the blastocysts. We drew a line with thickness ~1 μm along the cell-cell junctions between two mural TE cells, starting from the apical-most part of the tight junction towards the adherens junction. The mean intensity of the junctional or adherens markers were then plotted accordingly. For skeletal analysis, the continuity of signal for the di-phosphorylated (Thr18/Ser19) form of myosin regulatory light chain (ppMRLC) was analysed in mural TE of wild-type embryos fixed at E3.75 and E4.25. Phalloidin or ZO1 were co-stained as reference markers for junctional continuity. Maximum projections were performed for confocal slices containing junctions of mural TE cells. Thresholding and binarisation were applied before using the skeletonize function to extract the integrated density for ppMRLC and junctional marker. I_ppMRLC_/I_junction_ < 1 indicates punctate ppMRLC signal while I_ppMRLC_/I_junction_ ≈ 1 indicates a continuous linear signal.

#### Cell count

Cell count was performed using Modular Interactive Nuclear Segmentation (MINS-1) package running on MATLAB, as described in previous publication^37^. Multi channel images containing DAPI and Cdx2 channel were pre-processed in FIJI to reduce cytoplasmic or ZP noise prior to running MINS-1. ICM cell number was obtained as a difference between the total cell number (from DAPI) and the TE cell number (Cdx2+ nuclei), and counter-checked with manual count of Sox2^+^ and Gata4^+^ nuclei.

All statistical analysis was performed using the software Origin (version 8.5.6; OriginLab, Northampton, MA). Significances for data displaying normal distributions were calculated with unpaired Student’s *t*-test (two-tailed, unequal variance). All box plots extend from the 25^th^ to 75^th^ percentiles (horizontal box), with a line at the median and whiskers extending to max/min data points.

